# Anti-arthralgic activity of the n-hexane extract (HTp) of yellow oleander seeds; *Thevetia peruviana* (Pers.) K. Schum

**DOI:** 10.1101/2021.07.27.453963

**Authors:** Joy Ifunanya Odimegwu, Fatiha Oyebola Olabisi

**Affiliations:** Department of Pharmacognosy, Faculty of Pharmacy, College of Medicine Campus, University of Lagos, PMB 12003. Idiaraba, Lagos. Nigeria

**Keywords:** Arthralgia, Herbal medicine, Pain, *Thevetia peruviana*, yellow oleander

## Abstract

*Thevetia peruviana* (Pers.) K.Schum. (*Apocynaceae*) seeds are known to possess cardioactive glycosides such as thevetin A, thevetin B, nerifolin etc. They are also used locally for general pain relief for which there is no scientific evidence to our knowledge. Arthralgia is regarded generally as pain without inflammation. It is endemic in the society and sufferers continue to imbibe pain relieving drugs in their tons all over the world. Analgesic activity test was carried out using the formalin-induced pain models, at 0.1g, 0.2g and 0.3g/kg doses of n-hexane extracts of *Thevetia peruviana* seeds (HTp) in Wistar mice. Diclofenac was used as positive control. Acute toxicity test was carried out at doses of 1000, 2500 and 5000 mg/kg weight of test subject. It was observed that HTp at concentrations of 0.1g, 0.2g and 0.3g/kg showed significant analgesic effect at compared to the control. The percentage inhibition observed was 29.60%, 44.80% and 50.72% for the early pain phase and 100% for the late pain phase respectively, indicating HTp’s NSAID-like property. HTp showed the highest percentage inhibition at 300 mg/kg (50.72 %) and significant (P<0.005) pain reduction. HTp did not produce any toxicity up to a dose of 5000 mg/kg weight which is very interesting as the seeds are known for their toxicity due to the cardiac glycoside presence. The results of the study suggest that HTp does indeed relieve pain significantly in a dose dependent manner, thus justifying its use in management of arthralgia.

## 1.0 INTRODUCTION

Arthralgia from Greek originally literally means joint pain. But specifically those joint pains without inflammation. It therefore can be caused by a myriad diseases; arthritis, it could be a sign of of an allergic reaction, injury or infection. Most known pain killers have undesirable adverse effects; from acid reflux to ulcers. Plants have been a valuable source of new molecules and considered as an alternative strategy in search for new pain drugs with reduced adverse effects. Therefore many investigations on medicinal plants for their therapeutic purposes have increased (Silva *et al*., 2010).

Present research on plant species that have traditionally been used for the relief of the pain should still be seen as a logical research strategy in the search for new analgesic drugs (Silva, *et al*., 2010). The World Health Organization (WHO, 2004) had estimated that more than 80% of the world’s peoples rely on traditional medicine for their primary healthcare needs. The medicinal value of plants lies in some chemical substances that produce a definite physiological action on the human body. The most important of these bioactive compounds of plants are alkaloids, flavonoids, tannins and phenolic compounds (Edeoga *et al*., 2005).

*Thevetia peruviana* synonym of *Cascabela thevetia* (L.) Lippold.is known as a poisonous plant (The plant list, 2013). It is an evergreen tropical plant. Its fruit is green-black in color encasing a large seed.

**Fig. 1.**
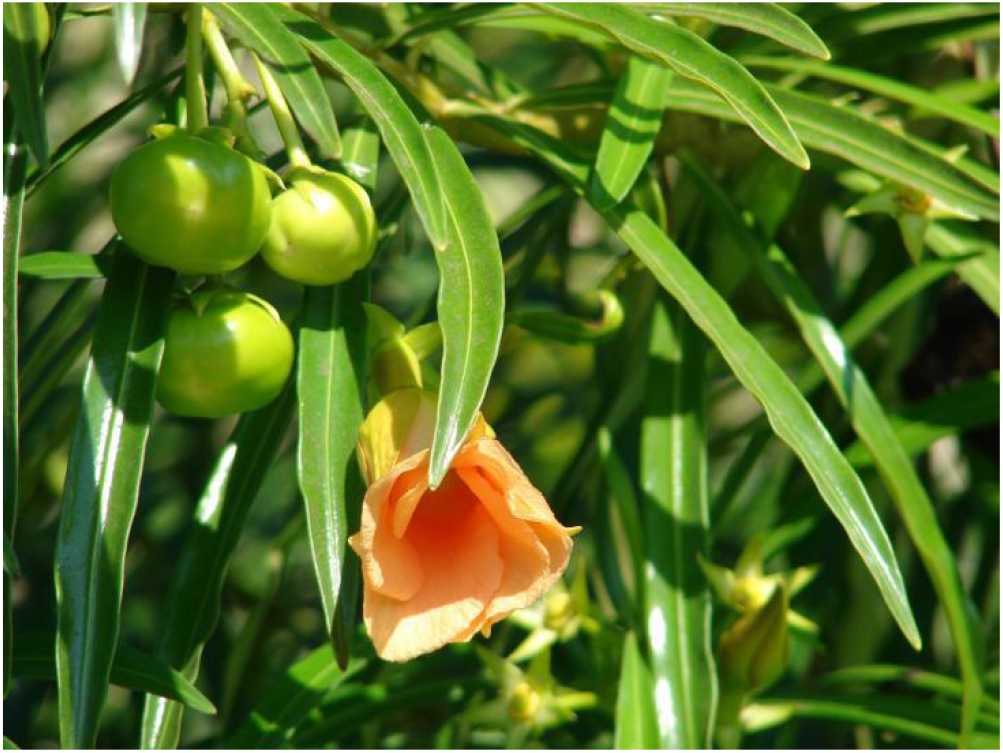
*Thevetia peruviana* plant with flower and seeds

### 1.1 The Problem of Pain

Pain is a common health problem with substantial socio-economic impact because of its high incidence (Julius and Basbaum, 2001 and Ballantyne, and Mao, 2003), it is a symptom characteristic of many diseases and a patho-physical response of living tissues to undesirable stimulus (Amol and Kallangouda, 2011). With many pathological conditions causing pain, tissue injury is viewed as the most immediate cause of pain and these results in the release of various chemical agents which are assumed to act on the nerve terminals either activating them directly or enhancing their sensitivity to other forms of stimulation (Mahesh, 2008; Kanodia, 2008).

Usually, pain is associated with discomfort, distress and perhaps agony, depending on its severity (Craig, 2007). It is an individual experience; it varies from one person to another as some people have higher threshold for pain than others. Pain is a subjective experience that can be perceived directly only by the sufferers. It involves the transduction of a noxious stimulus and also the cognitive and emotional process occurring in the brain (Vitor *et al*., 2008). Most pain resolves promptly once the painful stimulus is removed and the body has healed, but sometimes, it could persist even after the removal of the pain stimulus and healing of the body and sometimes rises in the absence of any detectable stimulus, damage or pathology (Quintner *et al*., 2015).

Pain signals travels from the periphery to the spinal cord along an A-delta or C-fibre. Because the A-delta fibre is thicker than the C fibre, and is thinly sheathed in an electrically insulating material (myelin), it carries its signal faster (5–30 m/s) than the unmyelinated C-fibre (0.5–2 m/s). Pain evoked by the A-delta fibres is described as sharp and is felt first. This is followed by a duller pain, often described as burning, carried by the C-fibres (Zagustin, 2013). These “first order” neurons enter the spinal cord via Lissauer’s tract (Skevington, 1995). Pain that is distinctly located also activates primary and secondary somatosensory cortex (Skevington, 1995).

Diclofenac, the standard used in this study is a nonsteroidal anti-inflammatory drug (NSAID) taken or applied to reduce inflammation and as an analgesic reducing pain in certain conditions. The primary mechanism responsible for its anti-inflammatory, antipyretic, and analgesic action is thought to be inhibition of prostaglandin synthesis by inhibition of cyclooxygenase (COX). Inhibition of COX also decreases prostaglandins in the epithelium of the stomach, making it more sensitive to corrosion by gastric acid. This is also the main side effect of Diclofenac which has a low to moderate preference to block the COX2-isoenzyme (approximately 10-fold) and is said to have, therefore, a somewhat lower incidence of gastrointestinal complaints than noted with indomethacin and aspirin (Dastidar *et al*., (2000). Diclofenac may also be a unique member of the NSAIDs. Some evidence indicates it inhibits the lipoxygenase pathways, thus reducing formation of the leukotrienes (also pro-inflammatory autacoids). It also may inhibit phospholipase A2 as part of its mechanism of action. These additional actions may explain its high potency - it is the most potent NSAID on a broad basis (Scholer, 1986).

The general aim of this study are to assess the anti-arthralgic activity of the extract of *Thevetia peruviana* seeds used locally for pain relief and also evaluate the phyto-components of HTp using GC/MS.

## 2.0 METHODOLOGY

### 2.1 PLANT COLLECTION AND IDENTIFICATION

*Thevetia peruviana* (yellow Oleander) Fruits were collected from a private garden. The plant material was identified, authenticated and assigned a voucher specimen number LUH: 7586 at the Herbarium of the Department of Botany, University of Lagos. Nigeria.

### 2.2 MATERIALS

N-Hexane, Tween 80, stopwatch, Formalin 1% (v/v)

### 2.3 SAMPLING AND SAMPLE PREPARATION

The seeds of *Thevetia peruviana* were oven dried and crushed using mortar and pestle. The crushed seeds were then pulverized with an electric blender. The powdered sample was stored at −4°C until required extraction.

### 2.4 EXTRACTION

This was carried out using a Soxhlet apparatus. The podered sample was immersed in analytical Hexane (100%) for 4-5 hours then filtered. The filtrate was concentrated and later left open for all hexane to evaporate.

### 2.5 EXPERIMENTAL ANIMALS

Male Wistar mice each weighing 20g were randomly selected. The animals were raised in the animal house of the Faculty of Pharmacology, University of Lagos, Nigeria. Animals were exposed to daily 12h light/12h dark cycle and free access to tap water and standard animal feeds. The animals were acclimatized in a well-ventilated room in the animal house under an ambient room temperature for 7 days prior to the experiment. The weights of the animals were taken before administration of the HTp and the animals separated in individual cages and marked to aid their identification. The effects of *Thevetia peruviana* was examined in the duration of paw licking in the formalin-induced pain model in the mice. All the ethical indications from the University of Lagos for animal handling were followed strictly.

### 2.6 ACUTE TOXICITY TEST

A single dose test was carried out. Ten mice were fasted for 16 hours and shared into groups; A-C Group A was administered 1000mg/kg, Group B were administered 2500mg/kg and Group C were administered 5000mg/kg of HTp. They were observed for 24 to 48 hours for any signs of behavioral changes and/or death.

### 2.7 EXPERIMENTAL PROCEDURE

Formalin-Induced hind-paw licking Test was performed using the method described by Hunskar and Hole, (1987) and as reported by Young *et al*. (2005) with some modifications. 25 mice were divided into five groups of five animals each for testing. The groups are as follows:

Group 1: Control - Normal saline (2ml/kg); Group 2: Standard Group - Diclofenac Sodium (9mg/kg)
Group 3: Test Sample 100mg/kg; Group 4: Test Sample 200mg/kg; Group 5: Test Sample 300mg/kg

The animals were fasted for 24 hours prior to the experiment. For the test sample, the HTp was administered in the form of a patch, but later, the volume of each administered dose was dropped at the back of the mice where it cannot be tampered. Each mice was kept in a separate cage.

## 3.0 RESULTS

### 3.1 ACUTE TOXICITY

The result of the acute toxicity of *T. peruviana* treatment with single doses of 1000mg/kg, 2500 mg/kg and 5000 mg/kg did not result in any toxic signs or mortality. This indicates that the LD50 is greater than 5000 mg/kg. This is interesting because this plant seed is known to be very toxic in water or alcohol, so this implies that HTp does not have the toxic principle.

### 3.2 GC/MS Analysis

The GC-MS analysis identified compounds from HTp and these were matched with those found in using the National Institute of Standard and Technology (NIST) Spectral Search Program Version 2.0.Database. The GC-MS chromatogram showed the twenty-six peaks of compounds detected. The results revealed that Oleic acid (50.36%), Palmitic acid (24.04%) and Stearic acid (12.30) were found to be the 3 major components in HTp. The rest were; Glycerin; Ethanone,1-(1H-pyrrol-2-yl)-; 2-Nonen-1-ol, (E)-; Octanoic acid; Naphthalene; 2-Tridecenal, (E)-; Dodecanoic acid; Diethyl Phthalate; Ar-tumerone; Curlone; Tetradecanoic acid; Hexadecanoic acid, methyl ester; 9-Hexadecenoic acid; Hexadecanoic acid, ethyl esther; 9,12-Octadecadienoic acid (Z,Z)-; 6-Octadecenoic acid, methyl ester, (Z)-; Hexadecanoic cid,1-(hydroxymethyl)-1,2-ethanediyl ester; 9-Octadecenoic acid 1,2,3-propanetriyl ester, (E,E,E)-; Hexadecanoic acid, 2-hydroxy-1-(hydroxymethyl) ethyl ester; Oleoyl chloride; Squalene; Nonacosane and Hexatriacontane.

### 3.3 FORMALIN TEST

HTp showed significant (P<0.005) analgesic effect at doses of 200 mg/kg and 300mg/kg in both early and late phase of the experiment compared to the control. In the early phase of the experiment, percentage inhibition of the paw licking was 44.80% and 50.72% while in the late phase the percentage inhibition was 100%. There is no significant difference between the dose of 100 mg/kg and the control at the both phases. Diclofenac showed the highest percent inhibition of 85.92% with little or no paw licking at early and later phases. There is also a significant (P<0.005) difference between the doses of HTp and the standard Diclofenac. As shown in Table 1.

**Table 1:**
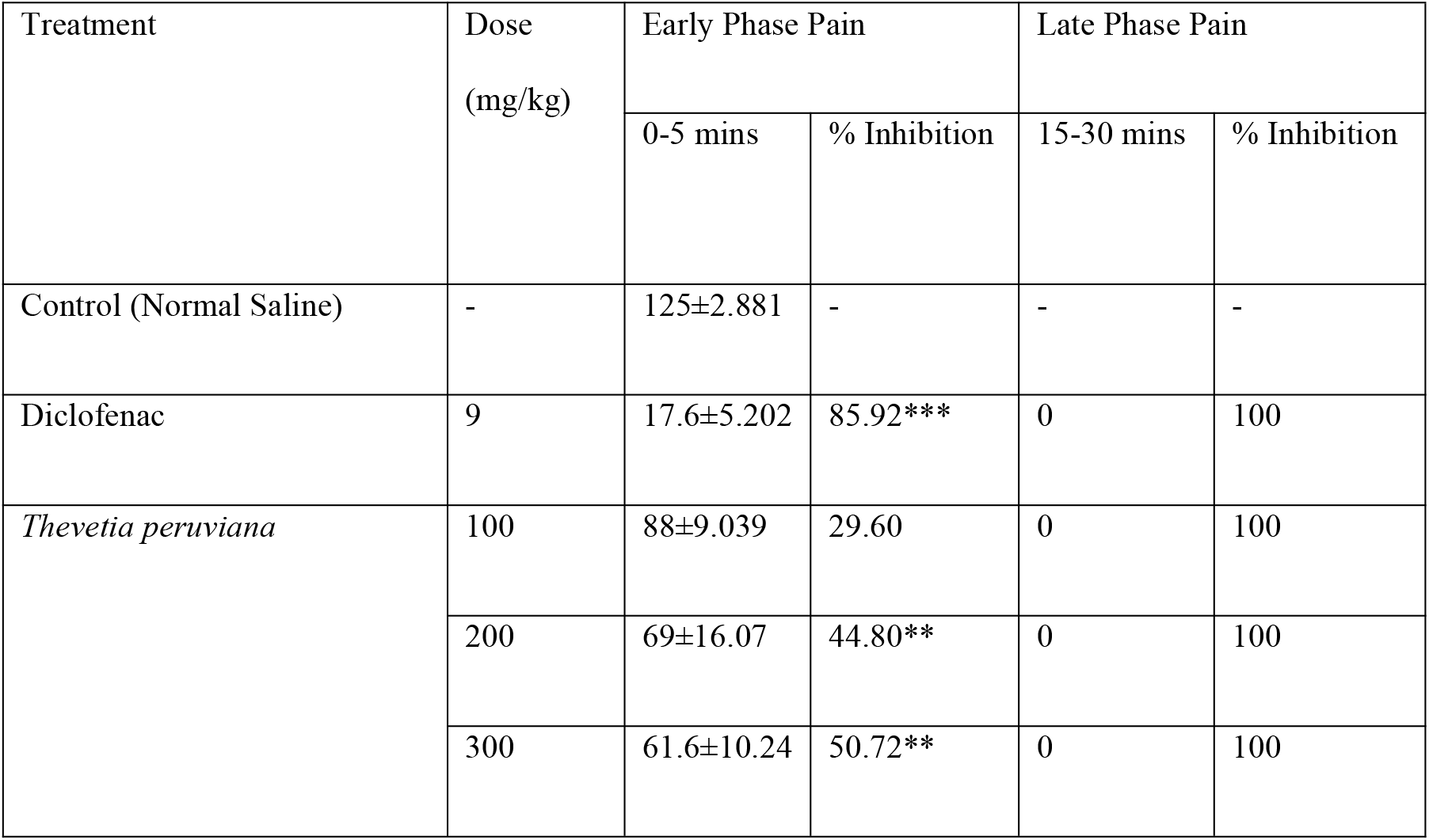
Analgesic effect of the hexane extract of *Thevetia peruviana* on formalin-induced pain model. Here, all values are expressed as Mean ± S.E.M (n=6) at ***P<0.005 very significant compared to control. Data was analysed by One-Way ANOVA

## 4.0 DISCUSSION

Oleic acid is known to possess anti-inflammatory activity (James *et al*., 1993). Palmitic acid has been reported to show anti-atherosclerotic properties (Sayeed *et al*., 2004). The presence of these bioactive constituents can further explain and validate the analgesic effects of HTp and thus, the anti-arthralgic actions of the extract. The selection of this model was informed by need to investigate the peripheral as well as the central mediated effect of the plant sample. It has been demonstrated previously that formalin test is believed to demonstrate the involvement of both central and peripheral pathways of pain (Ramirez *et al*., 2010).HTp demonstrated analgesic activity in blocking both phases of the formalin response. Manifestation in the early phase is due to the reduction in the neurogenic activity (Verma *et al*., 2005). This explains the function of HTp as analgesic and anti-inflammatory activities which mimics the action of NSAIDs. Although the effect of the formalin induced pain in the late phase was not observed in the control. Diclofenac was used as the standard drug. In relations to HTp, doses of 200mg/kg and 300mg/kg inhibited pain, although not as much as the standard, as there was a significant difference between the standard and HTp. This led to the inference that, higher doses are required to achieve a higher response of pain relief.

The effect of HTp on both phases suggest peripheral and central mechanism, but due to the fact that the control group did not show any paw licking even at phase two, it is difficult to conclude that HTp completely inhibited the pain at the late phase.

The result showed a dose dependent response, as the inhibition if pain was greatest in the 300mg/kg concentration of HTp, then the 200mg/kg and then the 100mg/kg. The graphical representation of the relationship between effects is shown in Fig. 2.

**Fig. 2:**
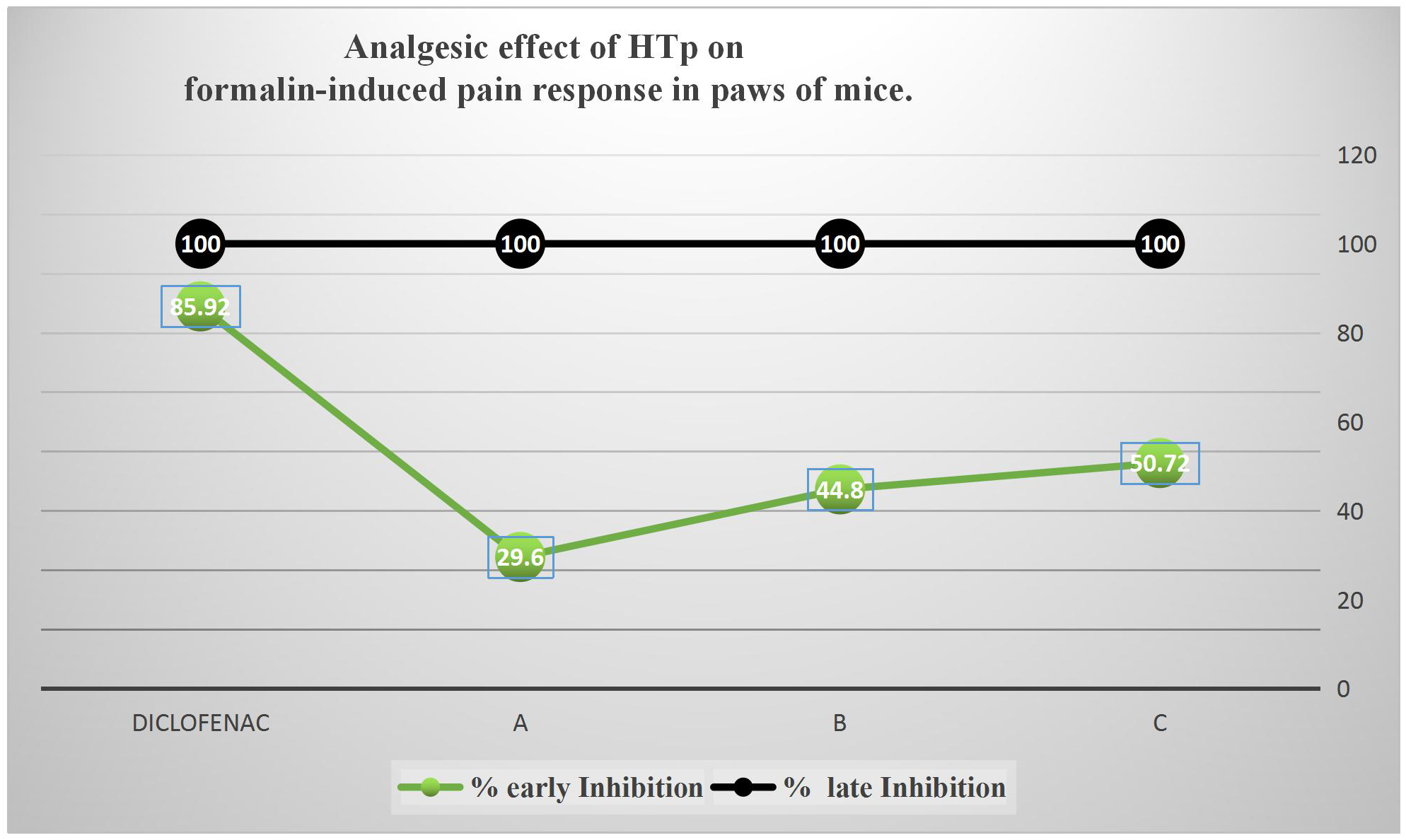
**R**elationship between anti-arthralgic activity of test samples and standard NSAID

Oral acute toxicity of *T. peruviana* on mice shows that no animal mortalities within 48 hours after treatment with HTp. Treatment with single doses of 1000mg/kg, 2500 mg/kg and 5000 mg/kg did not result in any toxic signs or mortality in the acute toxicity studies. This indicates that the LD50 is greater than 5000 mg/kg. This shows that HTp is safe at doses 5000mg/kg or less. This is interesting because this plant seed is known to be very toxic when eaten, The obtained result therefore implies that the n-hexane HTp is not toxic.

## CONCLUSION

The results from the study provide evidence for the anti-arthralgic activity of HTp, as confirmed by the Formalin-Induced Pain model which supports and validates its use in the local communities in the treatment of arthralgia.

